# BERTMeSH: Deep Contextual Representation Learning for Large-scale High-performance MeSH Indexing with Full Text

**DOI:** 10.1101/2020.07.04.187674

**Authors:** Ronghui You, Yuxuan Liu, Hiroshi Mamitsuka, Shanfeng Zhu

## Abstract

**Motivation:** With the rapid increase of biomedical articles, large-scale automatic Medical Subject Headings (MeSH) indexing has become increasingly important. FullMeSH, the only method for large-scale MeSH indexing with full text, suffers from three major drawbacks: FullMeSH 1) uses Learning To Rank (LTR), which is time-consuming, 2) can capture some pre-defined sections only in full text, and 3) ignores the whole MEDLINE database.

**Results:** We propose a computationally lighter, full-text and deep learning based MeSH indexing method, BERTMeSH, which is flexible for section organization in full text. BERTMeSH has two technologies: 1) the state-of-the-art pre-trained deep contextual representation, BERT (Bidirectional Encoder Representations from Transformers), which makes BERTMeSH capture deep semantics of full text. 2) a transfer learning strategy for using both full text in PubMed Central (PMC) and title and abstract (only and no full text) in MEDLINE, to take advantages of both. In our experiments, BERTMeSH was pre-trained with 3 million MEDLINE citations and trained on approximately 1.5 million full text in PMC. BERTMeSH outperformed various cutting edge baselines. For example, for 20K test articles of PMC, BERTMeSH achieved a Micro F-measure of 69.2%, which was 6.3% higher than FullMeSH with the difference being statistically significant. Also prediction of 20K test articles needed 5 minutes by BERTMeSH, while it took more than 10 hours by FullMeSH, proving the computational efficiency of BERTMeSH.

## 1 Introduction

As a comprehensive controlled vocabulary, Medical Subject Headings (MeSH) has been developed and maintained by the National Library of Medicine (NLM) for indexing, cataloging and searching of biomedical information (Sayers *et al*., 2020). As of 2020, there are 29,640 MeSH main headings (MHs)^1^. One of the most important usages of MeSH is to index the largest biomedical literature database, MEDLINE, which currently covers more than 5,200 journals and 26 million citations (Sayers *et al*., 2020). Currently each MEDLINE citation is annotated with 13 MHs on average, which can be utilized in many applications in biomedical text mining and information retrieval (Lu *et al*., 2009; Stokes *et al*., 2009; Gu *et al*., 2013; Huang *et al*., 2011; Zhu *et al*., 2009). Accurate MeSH indexing is thus crucial for biomedical researchers, who are generating new hypotheses and seeking to make new discoveries.

In 2019, 956,390 citations have been added into MEDLINE, which is around 5% increase over 2018 (904,636)^2^. The vast majority of these citations are manually indexed with MHs by human curators in NLM, with an average annotation cost of $9.4 per citation (Mork *et al*., 2013). To deal with the rapid growth of MEDLINE, NLM has developed a software tool, Medical Text Indexer (MTI), to facilitate the MeSH indexing task in automated and semi-automated modes (Aronson *et al*., 2004; Mork *et al*., 2017). Currently around 5% of MEDLINE citations are annotated automatically, where MTI provides MHs without human intervention^3^. On the other hand, around 18% of MEDLINE citations are annotated semi-automatically, where human curators review (and possibly revise) the MHs recommended by MTI. Note that MTIs use only the title and abstract of each citation to recommend MHs, while human curators in NLM check the full text to finish the MeSH indexing task. Meanwhile, the number of available full text in PubMed Central (PMC) reaches 5.9 million in Jan 2020 ^4^. With the rapid growth of full text biomedical articles, it is an imperative task to develop an accurate and efficient automatic MeSH indexing method for large-scale full text.

From a machine learning perspective, automatic MeSH indexing can be deemed as a large-scale multi-label learning problem, where MHs are labels, citations are instances, and each citation is associated with multiple MHs (Liu *et al*., 2015). To advance the performance of automatic MeSH indexing, many advanced machine learning methods have been developed to address this challenging problem in the last few years, such as MetaLabeler (Tsoumakas *et al*., 2013), MeSHNow (Mao and Lu, 2017), MeSHLabeler (Liu *et al*., 2015), DeepMeSH (Peng *et al*., 2016), AttentionMeSH (Jin *et al*., 2018), MeSHProbeNet (Xun *et al*., 2019) and FullMeSH (Dai *et al*., 2020). Different from all other methods using title and abstract only, FullMeSH makes use of full text to extract different sections, and utilizes Learning To Rank (LTR)(Li, 2011) to integrate the evidence generated from each section to improve the performance of MeSH indexing. However, FullMeSH suffers from three major drawbacks: 1) the performance of FullMeSH drops significantly if pre-defined sections are missed in the full text. This is because FullMeSH relies on pattern matching to extract five standard sections: *Title and Abstract, Introduction, Methods and Materials, Result and Experiment, Conclusion and Summary*. Many biomedical articles however do not have all these five sections. Additionally, pre-defined patterns can hardly deal with all kinds of variations of section names. 2) FullMeSH relies on LTR to integrate many different types of evidence generated from each section, which is complicated, laborious and time consuming. 3) although FullMeSH uses the full text of PMC open access data to train the model, FullMeSH cannot take advantage of the whole MEDLINE database.

In this work, we propose a novel deep learning based method, BERTMeSH, to improve the performance of large-scale MeSH indexing with full text. The main contributions of BERTMeSH are as follows:

- To the best of our knowledge, BERTMeSH is the first end-to-end deep learning based automatic MeSH indexing method for full text of large-scale biomedical documents (>1M) and all 29k MHs. In contrast to FullMeSH of LTR, BERTMeSH adopts a deep multi-label model with attention mechanism to capture the most relevant part of text for each label. In addition, we use Bidirectional Encoder Representations from Transformers (BERT) (Devlin *et al*., 2019) in BERTMeSH to encode the input text. In general natural language processing (NLP), we can find many successful applications of BERT, which can better capture text context and semantics in NLP. As BERTMeSH is an end-to-end deep model, it is more convenient for the developer to train, test, deploy and maintain the model.
- Without relying on pattern matching to distinguish different sections, BERTMeSH can robustly use the content of full text. Besides the title and abstract, the top four longest sections of each full text biomedical article are extracted as input in BERTMeSH. This avoids the missing section problem, caused by pattern matching and heterogeneous nature of different articles.
- In addition to utilizing full text in PMC, BERTMeSH can take advantage of the whole MEDLINE to improve the performance of large-scale MeSH indexing. The MEDLINE database is utilized in two distinct ways: 1) the whole MEDLINE has been used as corpus to train BERT model (BioBERT(Lee *et al*., 2020)), which can better reflect the characteristics of biomedical articles, and is used to encode text in BERTMeSH. 2) the network parameters in BERTMeSH are pre-trained by millions of MEDLINE citations, which greatly boost the performance.
- We conducted a thorough experiment for validating BERTMeSH by using PMC Open Access Subset (>1.4M) with 20,000 test articles. BERTMeSH achieved Micro F-measure of 69.2%, being 6.3% and 6.0% higher than those of the two start-of-the-art MeSH indexing methods, FullMeSH (65.1, trained on whole PMC Open Access Subset) and DeepMeSH (65.3, training on whole MEDLINE), respectively. Furthermore, BERTMeSH improves around 8.3% in Micro F-measure over FullMeSH for indexing full text articles with at least one missing section.

## 2 Related Work

### 2.1 Large-scale MeSH indexing based on title and abstract

To the best of our knowledge, all state-of-the-art large MeSH indexing methods, except FullMeSH, use title and abstract only. A classic method for large-scale MeSH indexing is NLM-developed MTI (Aronson *et al*., 2004; Mork *et al*., 2017) with two components: PubMed-Related citations (PRC) and MetaMap Indexing (MMI). PRC is a modified *k*-nearest neighbor (KNN) algorithm, to obtain the MHs of some most similar citations; MMI uses MetaMap to extract biomedical concepts from title and abstract, which are then mapped to MHs. These two sets of MHs are combined, ranked and recommended to the NLM curators after some post-processing, such as applying indexing rules.

Since 2013, many more advanced machine learning based methods have been proposed to tackle the problem of large-scale MeSH indexing, which is greatly facilitated by BioASQ challenges (2013-2019) that provide a practical and realistic benchmark for performance comparison (Tsatsaronis *et al*., 2015). Based on the machine learning techniques used, these automatic methods can be divided into three categories. (i) Binary relevance (BR); The best system in BioASQ 2013 developed by Tsoumakas *et al*. (2013), MetaLabeler, belongs to this category, where a linear SVM classifier is trained for each MH independently. Given a test citation, the candidate MHs are ranked according to the prediction score of each MH classifier. (ii) Learning to rank (LTR); MeSH Now (Mao and Lu, 2017), MeSHLabeler (Liu *et al*., 2015) and DeepMeSH (Peng *et al*., 2016) are three representative methods in this category. The main idea is to model MeSH indexing as a problem of ranking multiple MHs, where top ranked MHs are recommended as true labels. LTR has been successfully applied in the field of information retrieval, such as web searching. In the case of MeSH indexing, multiple evidence generated from different text representations and machine learning models are effectively integrated by LTR to improve the performance. Note that MeSHLabeler achieved the first place in BioASQ 2014 and 2015, while DeepMeSH achieved the first place in BioASQ 2016, 2017 and 2019. (iii) Deep Learning; AttentionMeSH (Jin *et al*., 2018) and MeSHProbeNet (Xun *et al*., 2019) are two recent deep learning based methods, which both use deep recursive neural network (RNN) and attention mechanism. Specifically, MeSHProebNet achieved the first place in BioASQ 2018 and the second place in BioASQ 2019, while AttentionMeSH achieved the third place in BioASQ 2018.

Note that all these methods used title and abstract only, which cannot take advantage of rich information in full text. In addition, although AttentionMeSH and MeSHProbNet are two deep learning based methods, they cannot enjoy the recent progress in pre-training in natural language processing (NLP).

### 2.2 BERT: Bidirectional Encoder Representations from Transformers

Deep learning models for NLP tasks used raw texts as inputs to capture rich semantic context information. Previously deep learning methods for NLP tasks usually used word embeddings (Mikolov *et al*., 2013) pre-trained on a large corpus to covert words to their corresponding dense semantic vectors, such as Word2Vec (Mikolov *et al*., 2013) and GloVe (Pennington *et al*., 2014). In spite of some successful applications in NLP including MeSH indexing (Peng *et al*., 2016), this type of word embedding uses an identical vector (representation) for the same word in different sentences, which cannot model the local context very well. Some pre-trained contextual text representations were then developed for replacing the single word embedding, such as ELMo (Embeddings from Language Models) (Peters *et al*., 2018) and BERT (Bidirectional Encoder Representation from Transformers) (Devlin *et al*., 2019). Different from previous language representation models like ELMo, BERT considers both left and right context of text when learning the language representation. The pre-trained BERT model has found many successful applications in NLP. Given the excitement about BERT, it was noted that it should be applied to biomedical NLP (Burns *et al*., 2019). Most recently, based on BERT, Lee *et al*. (2020) trained a biomedical domain specific language model BioBERT using biomedical text corpus such MEDLINE and PMC. They found that BioBERT improved the performance of several typical biomedical text mining tasks, such as biomedical name entity recognition, relation extraction and question answering. In this work, we used BioBERT for text representation, which greatly improves the performance of large-scale MeSH indexing.

## 3 Methods: BERTMeSH

### 3.1 Overview

Fig. 1 shows the architecture of BERTMeSH. For each biomedical article, we use the raw text from title and abstract and the *M*-1 longest sections from body text as our inputs. A pre-trained BERT layer (Devlin *et al*., 2019) is employed to obtain deep contextual representation of each word. For reducing the scale (the number of parameters) of our model, we use an identical BERT layer for all sections. Then we concatenate the outputs of all sections after the BERT layer as the representation of a given article. Following AttentionCNN, a deep component model of FullMeSH (Dai *et al*., 2020), we use a multi-label attention over the gained representation to capture the most relevant parts to each label, resulting in a different representation to each label. Finally, we use fully connected layers with sharing weights to obtain the predicted score to each label. To take advantage of both full text from PMC and a large amount of labeled citations from MEDLINE (which have title and abstract only), we use a transfer learning strategy. Specifically, BERTMeSH is pre-trained with millions of MEDLINE citations first, and then fine tuned with PMC full text data.

**Fig. 1.**
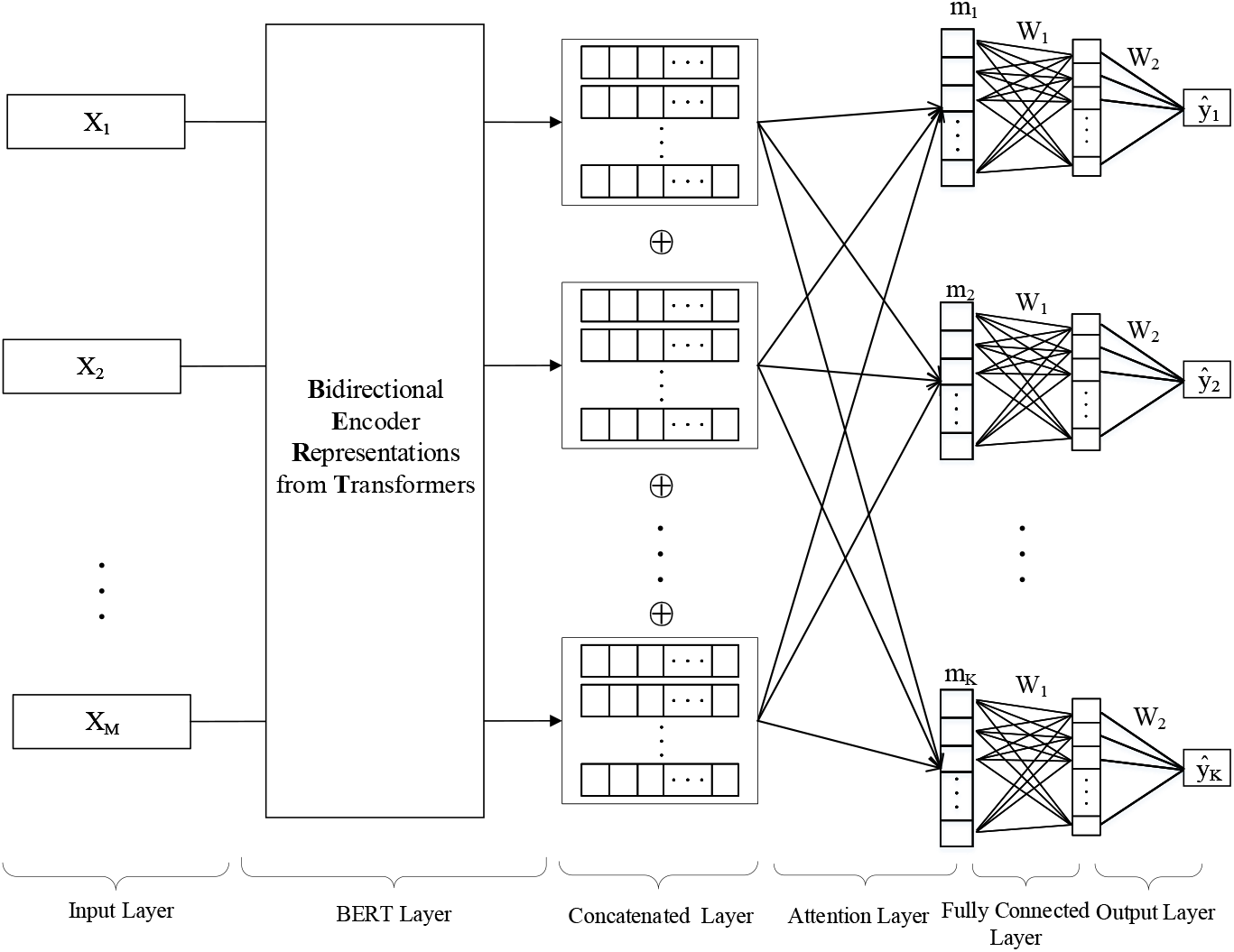
The architecture of BERTMeSH

### 3.2 Input Layer

For each sample, we use the raw text of *M* sections as our inputs, including the title and abstract section and the *M*-1 longest sections from the body text. The input ***X**_k_* of the *k*-th section for a given sample is as follows:

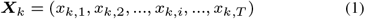

where the *x_k,i_* is the *i*-th word in the *k*-th section and *T* is the text length of each section.

### 3.3 BERT Layer

We use BioBERT (Lee *et al*., 2020) as our text representation model. BioBERT fine-tunes pre-trained BERT-base on MEDLINE and PMC corpus. The output ***H**_k_* of the *k*-th section is as follows:

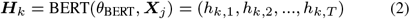

where 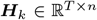, *n* is the hidden size of BERT, *θ*_BERT_ is the weight parameter of BERT Layer, and 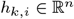 is the representation of the *i*-th word in the *k*-th section.

### 3.4 Concatenated Layer

For a given biomedical article, we concatenate *M* outputs of BERT layer over all *M* sections as follows:

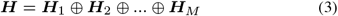

where 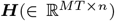 is the concatenated output, and *MT* = *M* × *T*. We denote 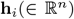 as the representation of the *i*-th word in the concatenated output ***H***.

### 3.5 Multi-label Attention

We use a multi-label attention to capture the most relevant part of text to each label to have different representations for each label. We use different attention parameters for each label. For the jth label, the attention we use is as follows:

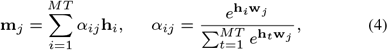

where **w**_*j*_ is the attention weight for the *j*-th label and **m**_*j*_ is the attention output for the *j*-th label.

### 3.6 Fully Connected Layer and Output Layer

BERTMeSH has one fully connected layer and one output layer. We set up that the fully connected layer and the output layer share the same parameter values for all labels, to emphasize the differences of attention among all labels and reduce the number of parameters. Finally, predicted probability *ŷ_j_* for the *j*-th label can be computed as follows:

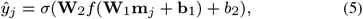

where 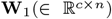 and 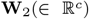 are parameters of the fully connected layer and output layer, respectively, and 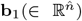 and 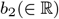 are bias terms, and *f* is a non-linear (activation) function.

### 3.7 Loss Function

BERTMeSH uses the binary cross-entropy loss, as the loss function, which is given as follows:

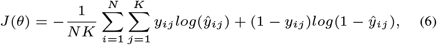

where *N* is the number of samples, *K* is the number of labels, *ŷ_ij_* ∈ [0,1] and *y_ij_* ∈ {0,1} are the predicted probability and true value, respectively, for the *i*-th sample and the *j*-th label.

### 3.8 Threshold for Each Label

After training, we compute the optimal threshold for each label over a threshold validation set, following (Pillai *et al*., 2013). We then select MHs with higher scores than the threshold as the final recommended MHs.

### 3.9 Pre-training with MEDLINE citations

Citations in PMC have full text, which has more useful information than title and abstract only. While the number of citations in MEDLINE with only title and abstract, is much larger than the number of citations in PMC. To improve the performance, we use transfer learning, to take advantage of both PMC and MEDLINE citations. Thus we first train BERTMeSH on MEDLINE citations with only title and abstract, and then fine-tunes this model with PMC citations.

## 4 Results

### 4.1 Data collection

We downloaded the whole PMC open access subset (by Oct. 2019)^5^ and obtained 3,221,713 citations. We also downloaded the whole MEDLINE collections (by Oct. 2019)^6^ and obtained 16,677,027 citations with abstract. For reducing the bias, we focused on manually indexed citations (not annotated by a “curated” or “auto” modes in MEDLINE) only in our work, which also have full text in PMC open access subset. Then we obtained a set of 1,495,063 PMC articles. Out of all these PMC articles, we used the latest 20,000 articles as the test set, another latest 200,000 articles except the test set as the threshold validation data set for all MHs, and the remaining 1.27M articles as the training set. We used the latest 3,000,000 MEDLINE citations as our MEDLINE pre-training dataset, which were annotated before the earliest date of the threshold validation data set. This means that this 3M MEDLINE citations have no overlap with our threshold validation and test datasets. Following the pattern matching method in FullMeSH, in addition to *Title and Abstract*, we extracted the 4 pre-defined sections from the test data, *Introduction, Methods and Materials, Result and Experiment, Conclusion and Summary*. Out of all 20,000 test articles, 13,641 (67.3%) articles had all these 5 sections, and the remaining 6,359 articles (32.7%) lost one or more sections in full text, where we call these two subsets the *complete subset* and *incomplete subset*, respectively.

### 4.2 Experimental Settings

As BERT Layer (with 12 transformer layer), we used BioBERT (Lee *et al*., 2020), which fine-tuned pre-trained BERT-base (Devlin *et al*., 2019) on MEDLINE and PMC open access subset with *n* = 768 and *T* = 512. We used the Adam optimizer (Kingma and Ba, 2014). Also we used a dropout with the drop rate of 0.5 and early stopping to avoid overfitting. We used five sections for training by PMC articles, including title and abstract, and the four longest sections (*M* = 5). There are six variants of BERTMeSH. First three variants use title and abstract only. They use PMC data, MEDLINE citations, and MEDLINE citations with Word2Vec and RNN instead of BERT layer, respectively. The other three variants use full text, and in practice, use PMC data only without pre-training, PMC data with pre-trained network by PMC abstracts, and PMC data with pre-trained network by MEDLINE citations.

Since the implementation of MeSHProbNet and AttentionMeSH for large-scale MeSH indexing is not available, we used DeepMeSH and FullMeSH as two competing methods, which are the state-of-the-art MeSH indexing methods using abstract and full text, respectively. Following the original paper (Dai *et al*., 2020), we implemented DeepMeSH and FullMeSH. Specifically, the same 20,000 latest PMC articles were used as the test set. Other 40,000 latest PMC articles were extracted, and half of the 40,000 articles were randomly chosen to train the ranking model, and the rest were used to train the model to predict the number of MHs annotated for a given article. Finally the remaining 1.4M articles were used as the training data. Note that FullMeSH used the full text of PMC, while DeepMeSH used the title and abstract only. In addition, we also checked the performance of DeepMeSH using the whole MEDLINE collection as the training data, after removing the above 60,000 citations.

### 4.3 Performance evaluation measures

Let *K* be the size of all labels (MHs), and *N* be the number of instances (citations). Let *y_i_* and *ŷ_i_* ∈ {0, 1}^*K*^ be the true and predicted labels for instance *i*, respectively. For performance evaluation, we used the three groups of most common metrics: Micro (precision (MiP), recall (MiR) and F-Measure (MiF)), Macro (precision (MaP), recall (MaP) and F-Measure (MaF)) and Example Based (precision (EBP), recall (EBR) and F-Measure (EBF)), which are defined as follows:

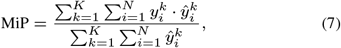

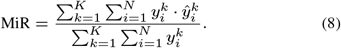

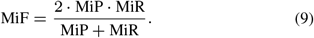

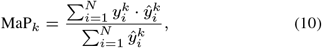

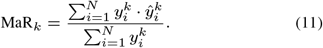

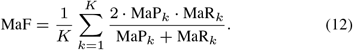

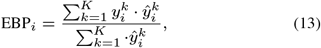

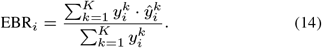

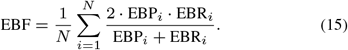

### 4.4 Experimental results

By using the 20,000 benchmark test articles, we compared the performance of BERTMeSH with DeepMeSH using title and abstract only, and then with FullMesH using full text. We mainly focused on MiF, the primary evaluation metric in the BioASQ challenge.

#### 4.4.1 Performance comparison with title and abstract only

Table 1 shows the performance comparison result of two settings of DeepMeSH (trained with PMC and trained with MEDLINE) and three settings of BERTMeSH (trained with PMC, trained with MEDLINE, and trained with MEDLINE using Word2Vec and RNN instead of BERT Layer). In Table 1, BERTMeSH outperformed DeepMeSH under all settings. BERTMeSH trained on MEDLINE achieved the best performance among the five settings. Specifically, BERTMeSH trained on MEDLINE achieved MiF of 0.678, being followed by BERTMeSH trained with PMC (0.667), BERTMeSH trained with MEDLINE using Word2Vec and RNN instead of BERT layer (0.662), DeepMeSH trained with MEDLINE (0.653), and DeepMeSH trained with PMC (0.639). We can see that with more training data (using MEDLINE instead of PMC), the performance of both DeepMeSH and BERTMesH was improved significantly in all three F-measures. For example, the MaF of DeepMeSH increased from 0.495 to 0.540. Another finding is that, without BERT layer, the performance of BERTMeSH trained with MEDLINE is even worse than BERTMeSH trained with PMC. For example, BERTMeSH trained with MEDLINE without BERT layer achieved MiF of 0.662, where BERTMeSH trained with PMC achieved MiF of 0.667. This highlights the power of deep contextual representation for improving the performance of MeSH indexing.

**Table 1.** Performance comparison of BERTMeSH and DeepMeSH by using title and abstract only.

| Method | Training | pre-trained | Full Text | MiF | MiP | MiR | MaF | MaP | MaR | EBF | EBP | EBR |
| --- | --- | --- | --- | --- | --- | --- | --- | --- | --- | --- | --- | --- |
| DeepMeSH | PMC | × | × | 0.639 | 0.669 | 0.612 | 0.495 | 0.633 | 0.502 | 0.631 | 0.667 | 0.627 |
| DeepMeSH | MEDLINE | × | × | 0.653 | 0.690 | 0.621 | 0.540 | 0.657 | 0.545 | 0.646 | 0.687 | 0.638 |
| BERTMeSH w.o. BERT Layer | MEDLINE | × | × | 0.662 | 0.695 | 0.631 | 0.525 | 0.668 | 0.526 | 0.652 | 0.700 | 0.643 |
| BERTMeSH | PMC | × | × | 0.667 | 0.696 | 0.640 | 0.512 | 0.663 | 0.517 | 0.657 | 0.700 | 0.650 |
| BERTMeSH | MEDLINE | × | × | **0.678** | **0.705** | **0.653** | **0.550** | **0.678** | **0.555** | **0.670** | **0.711** | **0.663** |

#### 4.4.2 Performance comparison with full text

Table 2 shows the performance comparison result of FullMeSH and three settings of BERTMeSH (pre-trained with PMC, pre-trained MEDLINE and without pre-training). In Table 2, BERTMeSH outperformed FullMeSH under all settings, and BERTMeSH pre-trained with MEDLINE achieved the best performance. Specifically, BERTMeSH pre-trained with MEDLINE achieved MiF of 0.692, which is 6.3% higher than FullMeSH (0.651). Even without pre-training, the performance of BERTMeSH reached MiF of 0.684, which is still much higher than the performance of FullMeSH. Note that if BERTMeSH is pre-trained with PMC itself, the performance increase is slight. For example, MiF ofBERTMeSHincreases from 0.684 to 0.685.

**Table 2.** Performance comparison of BERTMeSH and FullMeSH by using full text.

| Method | Training | pre-trained | Full Text | MiF | MiP | MiR | MaF | MaP | MaR | EBF | EBP | EBR |
| --- | --- | --- | --- | --- | --- | --- | --- | --- | --- | --- | --- | --- |
| FullMeSH | PMC | × | ✓ | 0.651 | 0.683 | 0.623 | 0.512 | 0.647 | 0.516 | 0.643 | 0.680 | 0.639 |
| BERTMeSH | PMC | × | ✓ | 0.684 | 0.711 | 0.660 | 0.526 | 0.666 | 0.532 | 0.674 | 0.715 | 0.667 |
| BERTMeSH | PMC | PMC | ✓ | 0.685 | 0.713 | 0.659 | 0.528 | 0.670 | 0.533 | 0.675 | 0.717 | 0.667 |
| BERTMeSH | PMC | MEDLINE | ✓ | **0.692** | **0.719** | **0.668** | **0.562** | **0.683** | **0.568** | **0.683** | **0.724** | **0.676** |

#### 4.4.3 Performance comparison with missing sections: validation on robustness

In Section 4.1, we divided the whole test data into two subsets: complete subset and incomplete subset. Again the complete subset consists of 13,461 (67.3%) test articles, each with all five sections, and the incomplete subset consists of 6,539 (32.7%) test article, each losing at least one section. In Table 2, BERTMeSH pre-trained with PMC had only little improvement, and then we examined the performance of three methods: BERTMeSH without pre-training, BERTMeSH pre-trained with MEDLINE, and FullMeSH. In addition, DeepMeSH using PMC abstracts was also examined as a baseline. Tables 3 and 4 show the performance results of all competing methods over the complete and incomplete subsets, respectively. BERTMeSH pre-trained with MEDLINE achieved the best performance in both cases, being followed by BERTMeSH without pre-training, FullMeSH and DeepMeSH. For example, over the complete subset, BERTMeSH pre-trained with MEDLINE achieved MiF of0.698, which wasfollowedby BERTMeSH without pre-training (0.691), FullMeSH (0.661) and DeepMeSH (0.646). Another point of note is that BERTMeSH is more robust than FullMeSH regarding missing sections, which can be seen from two viewpoints: 1) with missing sections, the MiF of BERTMeSH pre-trained with MEDLINE decreased 2.9%, i.e. from 0.698 to 0.678, and also that of BERTMeSH without pre-training decreased 3.6%, i.e. from 0.691 to 0.666. However, the decrease of FullMeSH was 5.3%, i.e. from 0.661 to 0.626. 2) over the complete subset, the MiF of BERTMeSH was 5.6% higher than that of FullMeSH, while over the incomplete subset, the MiF of BERTMeSH was 8.3% higher than that of FullMeSH. All these suggest BERTMeSH is more robust than FullMeSH with respect to the organization of sections in full text.

**Table 3.** Performance over full text with full sections.

| Method | Training | pre-trained | Full Text | MiF | MiP | MiR | MaF | MaP | MaR | EBF | EBP | EBR |
| --- | --- | --- | --- | --- | --- | --- | --- | --- | --- | --- | --- | --- |
| DeepMeSH | PMC | × | × | 0.646 | 0.679 | 0.616 | 0.501 | 0.637 | 0.507 | 0.639 | 0.677 | 0.630 |
| FullMeSH | PMC | × | ✓ | 0.661 | 0.705 | 0.623 | 0.516 | 0.662 | 0.516 | 0.654 | 0.703 | 0.637 |
| BERTMeSH | PMC | × | ✓ | 0.691 | 0.714 | 0.669 | 0.533 | 0.666 | 0.541 | 0.681 | 0.718 | 0.676 |
| BERTMeSH | PMC | MEDLINE | ✓ | **0.698** | **0.721** | **0.676** | **0.563** | **0.680** | **0.572** | **0.688** | **0.725** | **0.684** |

#### 4.4.4 Statistical performance superiority confirmation

By using the test set of 20,000 articles, we repeated boostrap with replacement 100 times, to generate 100 data sets. We then conducted paired *t*-test over 100 trials to examine the statistical significance on performance improvement between BERTMeSH and two state-of-the-art competing methods (DeepMeSH and FullMeSH). For BERTMeSH, we consider the two best settings: BERTMeSH with MEDLINE pre-training and BERTMeSH trained with MEDLINE. For DeepMeSH, hereafter we consider its best setting, which was trained with MEDLINE abstracts. Table 5 reports the predictive and statistical results of BERTMeSH, being compared with DeepMeSH and FullMeSH. In this table, below the performance values, the corresponding *p*-values are shown. Regarding the performance, BERTMeSH with PMC full text pre-trained with MEDLINE achieved the highest MiF of 0.692, being followed by BERTMeSH with MEDLINE abstract (0.678), DeepMeSH (0.653) and FullMeSH (0.651). Also from the *p*-values, which are far smaller than the regular statistical significance level, such as 0.05, the performance improvement by BERTMeSH was statistically significant. Overall the experimental results can be summarized into the following three points: 1) BERTMeSH outperformed other baselines for large-scale MeSH indexing, being statistically significant; 2) the performance of BERTMeSH was improved with full text; and 3) BERTMeSH shows the robustness over FullMeSH regarding missing sections in full text.

**Table 4.** Performance over full text with missing sections.

| Method | Training | pre-trained | Full Text | MiF | MiP | MiR | MaF | MaP | MaR | EBF | EBP | EBR |
| --- | --- | --- | --- | --- | --- | --- | --- | --- | --- | --- | --- | --- |
| DeepMeSH | PMC | × | × | 0.621 | 0.644 | 0.601 | 0.494 | 0.613 | 0.506 | 0.616 | 0.647 | 0.621 |
| FullMeSH | PMC | × | ✓ | 0.626 | 0.628 | 0.625 | 0.512 | 0.604 | 0.531 | 0.620 | 0.633 | 0.645 |
| BERTMeSH | PMC | × | ✓ | 0.666 | 0.700 | 0.634 | 0.528 | 0.656 | 0.535 | 0.659 | 0.709 | 0.648 |
| BERTMeSH | PMC | MEDLINE | ✓ | **0.678** | **0.714** | **0.646** | **0.559** | **0.677** | **0.564** | **0.672** | **0.722** | **0.661** |

**Table 5.** Statistical significance test by bootstrapping

| Method | Training | pre-trained | Full Text | MiF | MiP | MiR | MaF | MaP | MaR | EBF | EBP | EBR |
| --- | --- | --- | --- | --- | --- | --- | --- | --- | --- | --- | --- | --- |
| DeepMeSH | MEDLINE | × | × | 0.653 | 0.690 | 0.621 | 0.539 | 0.656 | 0.549 | 0.646 | 0.687 | 0.638 |
|  |  |  |  | 1.91e-169 | 7.66e-144 | 3.56e-168 | 3.92e-109 | 3.62e-110 | 2.96e-110 | 3.20e-164 | 1.65e-150 | 4.04e-159 |
| FullMeSH | PMC | × | ✓ | 0.651 | 0.683 | 0.623 | 0.513 | 0.645 | 0.522 | 0.643 | 0.680 | 0.639 |
|  |  |  |  | 1.48e-176 | 4.35e-16 | 2.41e-169 | 2.43e-144 | 1.38e-124 | 3.92e-143 | 1.32e-169 | 5.14e-160 | 7.84e-158 |
| BERTMeSH | MEDLINE | × | × | 0.678 | 0.705 | 0.653 | 0.551 | 0.677 | 0.562 | 0.670 | 0.711 | 0.663 |
|  |  |  |  | 3.58e-142 | 7.23e-129 | 1.83e-136 | 1.72e-97 | 1.06e-51 | 2.26e-98 | 9.39e-136 | 8.18e-127 | 1.44e-128 |
| BERTMeSH | PMC | MEDLINE | ✓ | **0.692** | **0.719** | **0.668** | **0.561** | **0.681** | **0.572** | **0.683** | **0.724** | **0.676** |

### 4.5 Result Analysis

#### 4.5.1 Performance comparison of three methods under different frequencies of MHs and articles

We divided MHs into five groups by the number of occurrences in training set: [0, 100), [100, 500), [500, 1,000), [1,000, 5,000) and [5000,). Fig. 2(a) shows the distributions of MHs and MHs indexing, when we divided training set into the above five groups. Fig. 2(b) shows the performance (average MaF) of BERTMeSH, FullMeSH and DeepMeSH, in each of the above five groups of MHs. For all groups, BERTMeSH achieved the best performance, being followed by DeepMeSH and FullMeSH. This highlights the advantage of BERTMeSH over DeepMeSH and FullMeSH, regardless of the frequency of MHs. Fig. 2(c) shows the distributions of the three methods regarding the best predictive methods (allowing ties) for each MH, for each of the five frequency groups. From Figs. 2(b) and 2(c), BERTMeSH outperformed DeepMeSH and FullMeSH, particularly more significantly for higher frequency groups. Another point of note is that for the most infrequent group, MaF of FullMeSH is much worse than DeepMeSH, while for the most frequent group, MaF of FullMeSH is only slight lower than DeepMeSH. This might be because: the size of the training set (MEDLINE) of DeepMeSH (which uses title and abstract only) is much larger than that (PMC) of FullMeSH (which allows to use all full text). That is, DeepMeSH would have used much more positive samples than FullMeSH. This might be very helpful for predicting infrequent MHs.

**Fig. 2.**
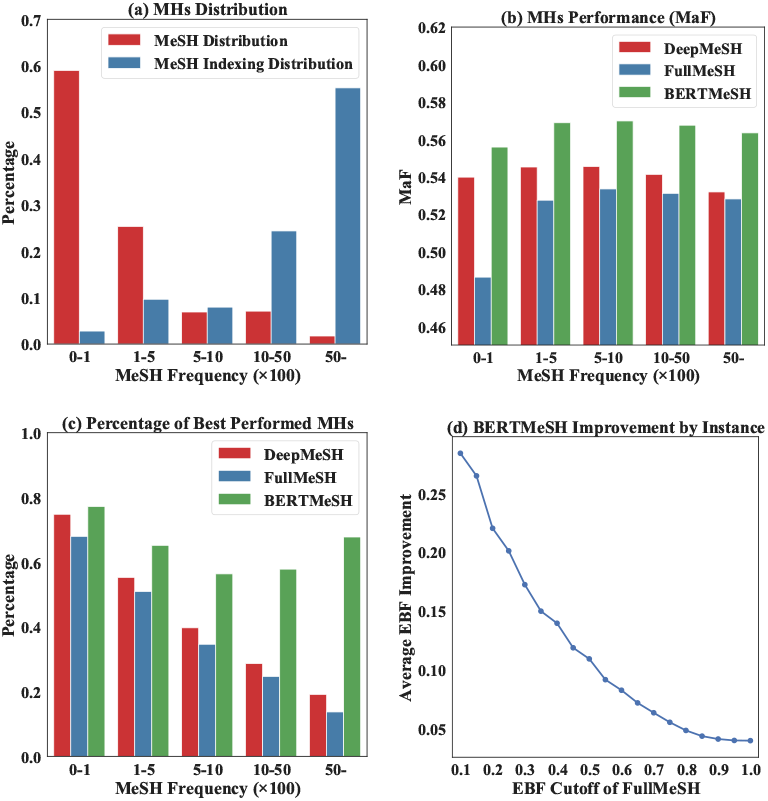
Performance comparison of DeepMeSH, FullMeSH and BERTMeSH

Furthermore, we compared the performance of BERTMeSH with FullMeSH with respect to instances (articles). We changed the size of test data, using EBF of each article by FullMeSH as the cut-off value, and then checked the performance improvement in the average EBF by changing the cut-off value. Fig. 2(d) shows the average EBF improvement of BERTMeSH over FullMeSH. From this figure, the improvement of BERTMeSH becomes larger, as EBF of FullMeSH became smaller, meaning that the article for which prediction is hard by FullMeSH can be better predicted by BERTMeSH.

#### 4.5.2 Performance comparison of different data usage for BERTMeSH under different frequencies of MHs and articles

We consider three data usages for BERTMeSH: 1) MEDLINE: BERTMeSH trained on only title and abstract in MEDLINE, 2) PMC: BERTMeSH trained on full text in PMC and 3) PMC+MEDLINE: BERTMeSH trained on full text in PMC with pre-training on MEDLINE. Fig. 3(a) shows the performance (MaF) of these three cases. First, focusing on MEDLINE and PMC only, MEDLINE outperformed PMC for all groups, except the most frequent group, particularly the performance advantage being wider for the groups with lower frequencies. This highlights the advantage of using MEDLINE for predicting less frequent MHs, which would be a large portion of the data set. On the other hand, PMC made better prediction on the high frequency group, although PMC had less training data (#articles would be smaller). For these most frequent MHs, which are usually important and general biomedical concepts, related information may be described in the full text other than abstract. In this case, PMC would be more advantageous than MEDLINE. Also Fig. 3(b) shows the distributions of the three data usages regarding the best predictive usage for each MH, for each of the five frequency groups. From Figs. 3(a) and 3(b), PMC+MEDLINE always outperformed the other two usages. This demonstrates that PMC+MEDLINE can enjoy the advantages of both pre-training by MEDLINE and training by full text of PMC.

**Fig. 3.**
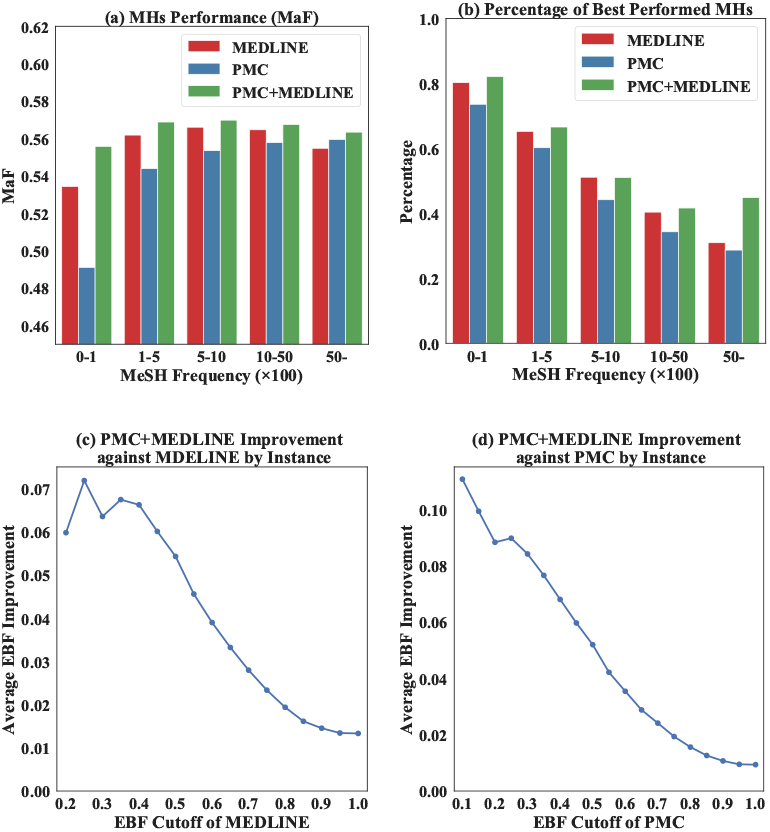
Performance comparison of BERTMeSH trained on MEDLINE, PMC and PMC with a pre-training over MEDLINE.

Furthermore, we again changed the size of test data, using the EBF of each article by MEDLINE (or PMC) as the cut-off value, and examined the performance improvement by PMC+MEDLINE. Figs. 3(c) and 3(d) show the average EBF improvement of PMC+MEDLINE over MEDLINE and PMC, respectively. The improvement of PMC+MEDLINE becomes larger, as the EBF of MEDLINE (or PMC) became smaller, meaning that the article for which prediction is hard by MEDLINE (or PMC) only can be better predicted by PMC+MEDLINE.

#### 4.5.3 Performance comparison of data usages on high frequency MHs by using Check Tags

Check Tags are a set of most frequent MHs, such as human, male, female and animal, meaning that Check Tags are likely to be mentioned in each article. In Section 4.5.2, we found that PMC (trained by full text) performed well for the most frequent MHs. We further check a similar but different setting of BERTMeSH by using Check Tags. We found 19 Check Tags that occur more than 150 times in our test set. Table 6 shows the F1-score of BERTMeSH by using three different data usages: MEDLINE, PMC and PMC+MEDLINE (which are the same as mentioned in Section 4.5.2) on the 19 Check Tags. In this table, PMC+MEDLINE achieved the highest F1-score in 14 out of 19 Check Tags, being followed by PMC, which achieved the highest in 6 Check Tags. Also both PMC and PMC+MEDLINE achieved the same average F1-score of 0.841. This result also suggests that the full text of PMC is useful for BERTMeSH to achieve good performance for highly frequent MHs.

**Table 6.**
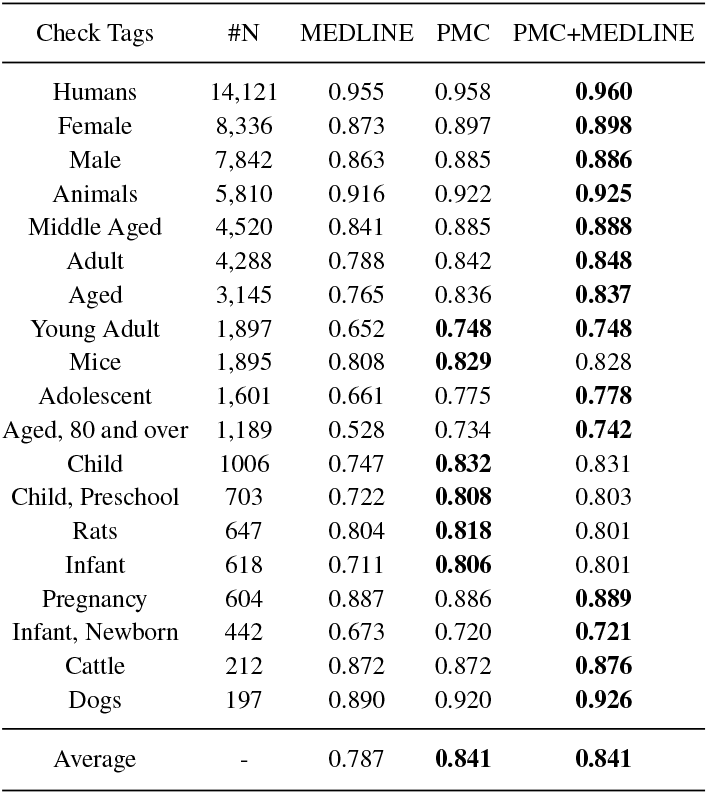
Performance (F1-score) comparison over Check Tags in test set (number of occurrence more than 150).

#### 4.5.4 Case Study

Table 7 shows the results of a sample article with PMID=31261512 (PMCID=PMC6616313). The first five rows of Table 7 are PMID (PMCID), title, abstract, part of main text, and MHs. The last four rows are the predicted MHs by DeepMeSH, FullMesH, BERTMeSH (MEDLINE) and BERTMeSH (PMC+MEDLINE), where correctly predicted MHs are highlighted in bold. Among the four methods, BERTMeSH (PMC+MEDLINE) achieved the highest F-measure of 0.947, while F-measure of the other three methods were all around 0.75. Specifically, three MHs, “Aged, 80 and over”, “ROC Curve” and “Young Adult” (all colored by red) were all correctly predicted by BERTMeSH (PMC+MEDLINE) but not by all other methods. We found that the descriptions most related with these three MHs cannot be seen in abstract, but appear in body text (in the Result section). For example, “The mean patient age was 55.4±14.1 years with a range of 18 to 91 years;” in the Result section implies two MHs: “Aged, 80 and over” and “Young Adult”. BERTMeSH could capture this information, while the other three methods could not use full text or could not capture the above semantics from full text efficiently. Overall this case study demonstrates the advantage of BERTMeSH over DeepMeSH and FullMeSH.

**Table 7.**
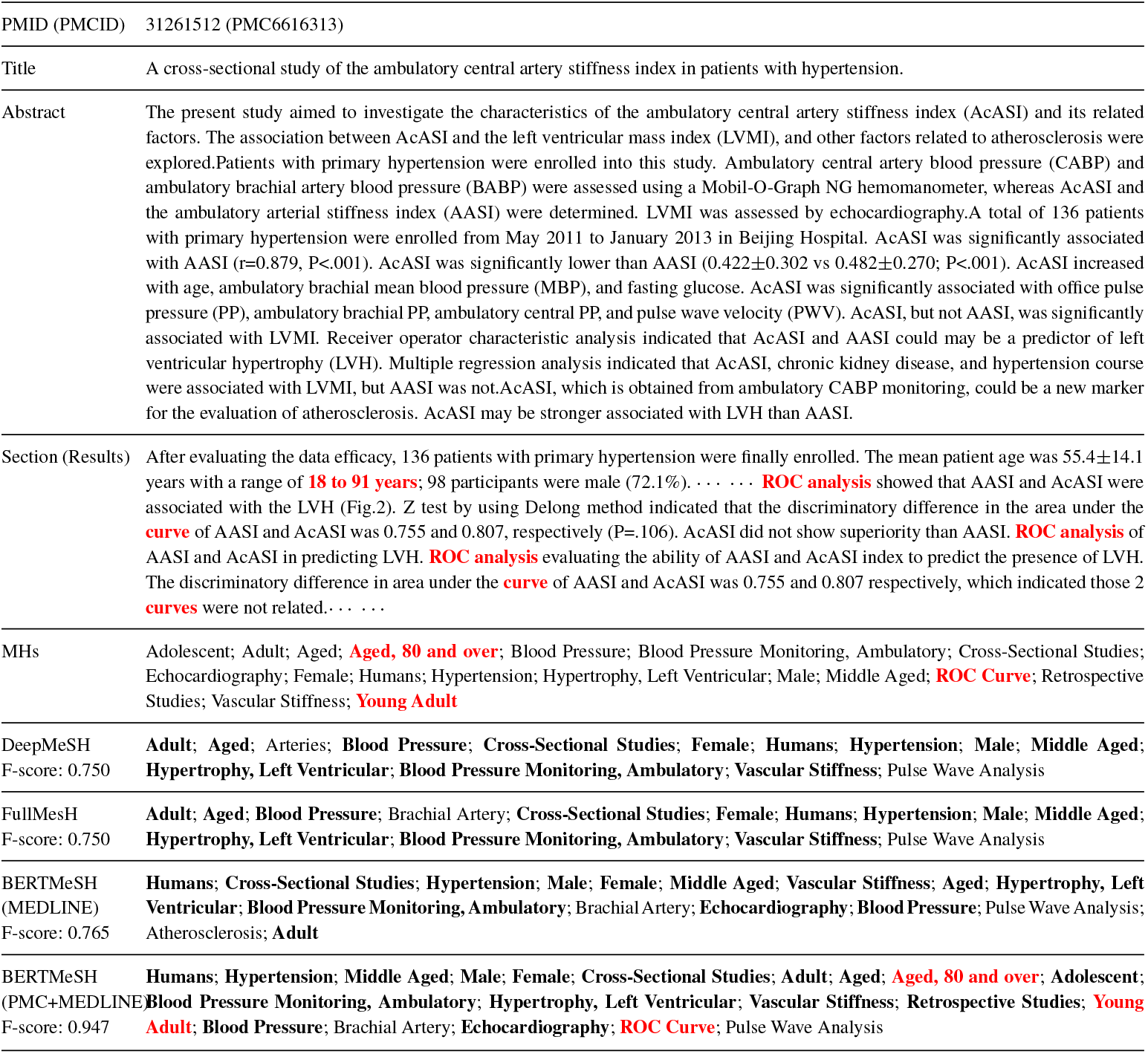
Case study for BERTMeSH.

### 4.6 Computation time

In our experiments, we used a server with 2 Intel Xeon E5-2678 V3 2.5GHz CPUs, 256G memory and 8 NVIDIA GTX 1080TI GPUs. Training BERTMeSH needed around 4 days, including pre-training, while FullMeSH needed around 7 days with a cluster server of **six** nodes, each being equipped with 128 GB RAM and two Intel XEON E5-4650 CPUs. Prediction by BERTMeSH needed around 5 minutes for 20,000 articles, while FullMeSH needed over 10 hours for 20,000 articles, mainly due to high computational cost of *k*-nearest neighbors.

## 5 Discussion and Conclusion

With the rapid growth of biomedical articles, automatic MeSH indexing with full text is becoming increasingly important. FullMeSH, only method of using full text for large-scale MeSH indexing had two serious problems: FullMeSH 1) uses LTR, which is laborious and time consuming, 2) can use only pre-determined sections in full text, limiting the advantage of using full text, and 3) ignores the whole MEDLINE database. To address these challenges, we have developed a computationally lighter model, BERTMeSH, which is flexible in section organization of full text, by using the state-of-the-art, deep contextual representation, BERT. Also BERTMeSH has pre-training, which allows BERTMeSH to use both MEDLINE (with only title and abstract) and PMC (with full text). Extensive experiments using 20K full text articles of PMC showed the efficiency, effectiveness and robustness of BERTMeSH over recent cutting-edge baselines. Interesting future work is to incorporate other recent techniques developed in the deep learning community to capture the interaction between different sections of the full text.

## Footnotes

1 https://www.nlm.nih.gov/databases/download/mesh.html

2 https://www.nlm.nih.gov/bsd/medline_pubmed_production_stats.html

3 https://www.nlm.nih.gov/pubs/techbull/ja18/ja18_indexing_method.html

4 https://www.ncbi.nlm.nih.gov/pmc

5 ftp://ftp.ncbi.nlm.nih.gov/pub/pmc

6 ftp://ftp.ncbi.nlm.nih.gov/pubmed/baseline

